# RNA-sequencing reveals a gene expression signature in skeletal muscle of a mouse model of age-associated post-operative functional decline

**DOI:** 10.1101/2021.10.05.461928

**Authors:** Samantha L. Asche-Godin, Zachary A. Graham, Adina Israel, Lauren M. Harlow, Weihua Huang, Zhiying Wang, Marco Brotto, Charles Mobbs, Christopher P. Cardozo, Fred C. Ko

## Abstract

This study aimed to characterize the effects of laparotomy on post-operative physical function and skeletal muscle gene expression in C57BL/6N mice at 3, 20 and 24 months of age to investigate late-life vulnerability and resiliency to acute surgical stress. Pre- and post-operative physical functioning were assessed by forelimb grip strength and motor coordination. Laparotomy induced an age-associated post-operative decline in forelimb grip strength that was greatest in the oldest mice. In contrast, while motor coordination declined with increasing age at baseline, it was unaffected by laparotomy. Moreover, baseline physical function as stratified by motor coordination performance (low vs. high functioning) in 24-month-old mice did not differentially affect post-laparotomy reduction in grip strength. RNA sequencing of soleus muscles showed that laparotomy induced age-associated differential gene expression and canonical pathway activation with the greatest effects in the youngest mice. Examples of such age-associated, metabolically important pathways that were only activated in the youngest mice after laparotomy included oxidative phosphorylation and NRF2-mediated oxidative stress response. Analysis of lipid mediators in serum and gastrocnemius muscle showed alterations in profiles of these mediators during aging and confirmed an association between such changes and functional status in gastrocnemius muscle. These findings demonstrate a mouse model of laparotomy which recapitulated some features of post-operative skeletal muscle decline in older adults following surgery, and identified age-associated, laparotomy-induced molecular signatures in skeletal muscles. Future research can build upon this mouse model to study molecular mechanisms of late-life vulnerability to acute surgical stress and resiliency to counter surgery-induced physical decline.

## Introduction

Surgery in older adults is associated with increased morbidity, mortality and healthcare costs compared to younger individuals.[1] In the United States, more than 50% of all surgeries are performed in adults >65 years of age. By one estimate, 7% of patients who underwent non-cardiac surgery experienced post-operative complications that led to a 78% increase in hospital cost and 114% increase in hospital length of stay.[2] Post-operative complications were more common with increasing age, with 20% of those greater than 80 years old experiencing one or more complications.[3, 4] Given that post-operative complications are common and have significant medical and financial consequences, and that surgeries are frequently performed in older adults, there is an urgent need to elucidate biological processes that increase risk for poor surgical outcomes in this vulnerable population.

It has been known for decades that surgery, including abdominal procedures, precipitates muscle weakness, weight loss, and inactivity in the post-operative convalescent period. In older adults, surgery induces muscle catabolism leading to muscle wasting [5], decreases limb muscle endurance and strength [6], delays recovery of muscle function [7], and causes persistent fatigue [8] and weight loss [9]. These adverse post-operative responses are associated with poor outcomes such as delayed return of physical activity, impaired recovery, and increased hospital readmission and mortality.[10-13] While the phenomenon of surgery-induced muscle and functional decline in older adults is widely recognized, its molecular underpinnings remain poorly understood. To date, few studies have investigated the molecular mechanisms regulating alterations in post-operative muscle function and physical decline in older individuals. Thus, there remains a gap in knowledge regarding the biology that mediates surgery-induced muscle and functional decline in older adults. As a first step to better understand this biology, we previously modeled surgical trauma introduced via laparotomy in 3-month-old male C57BL/6J mice and showed that this procedure induces coordinated, post-operative blood cell expression of unique inflammatory and oxidative or metabolic stress genes.[14] Moreover, in a model of post-operative stress in adult rats, changes in lipid mediators (LMs) that modulate inflammation in intestinal mucosa after laparotomy were previously observed.[15] In the current study, we sought to further characterize the effects of laparotomy on post-operative physical function, skeletal muscle gene expression and LM levels in C57BL/6N mice across the life span (3, 20 or 24 months of age) in order to investigate the impact of age and function on late-life vulnerability and resiliency to acute surgical stress.

## Results

### Laparotomy caused an age-associated post-operative decline in forelimb grip strength

C57BL/6N mice of different ages (3, 20 or 24-month-old) were trained to perform a forelimb grip strength task and then subsets of the animals underwent a laparotomy procedure or were observed as non-surgical controls. The general timeline of behavioral tests can be found in (Figure 1A). The average forelimb grip strength was lower in mice that underwent laparotomy compared to the control mice for all age groups on post-operative day (POD) 1 and 3 (p<0.0007). (Figure 1B and 1C) Moreover, on POD 3, the average grip strength for the 24-month-old laparotomy group was 21.5% lower than that of the 3-month-old laparotomy group (p<0.0007) and 19.8% lower than the 20-month-old laparotomy group (p=0.003), suggesting that the laparotomy procedure had the greatest adverse effect on the grip strength of 24-month-old mice.

**Figure 1:**
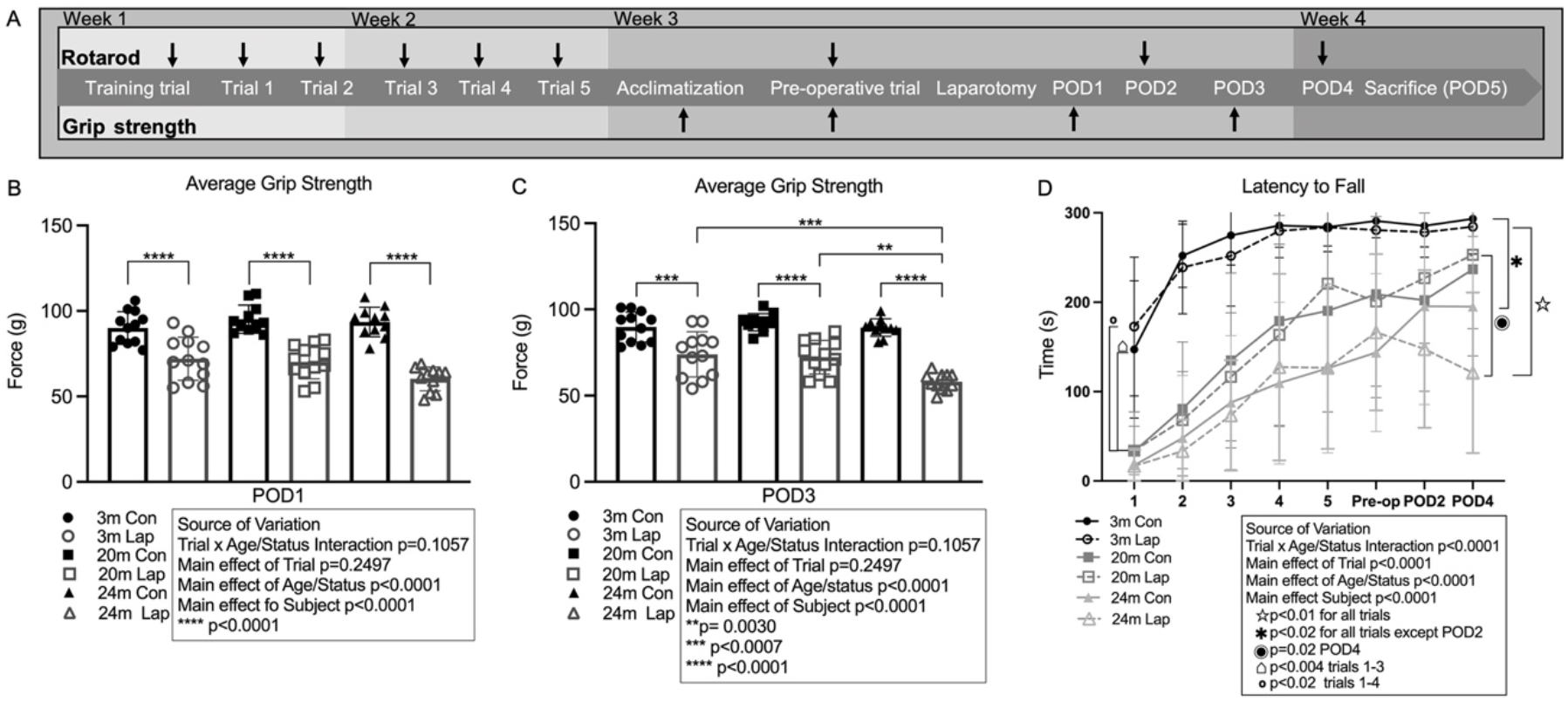
Behavioral analysis of 3-month, 20-month and 24-month-old C57BL/6N mice before and after laparotomy. (A) Timeline of behavioral testing. (B and C) Grip strength: the average forelimb grip strength force in grams was measured on POD 1 and 3. (D) Rotarod analysis: the latency to fall time was measured in seconds on the accelerating rotarod. N=11-12/group; error bars denote standard deviation. Statistical analysis performed by 2-way ANOVA with Tukey test for multiple comparisons; a repeated measures design was used for rotarod analysis. POD=post-operative day; Con=control; Lap=laparotomy.

### Motor coordination decreased with age but was not affected by laparotomy

In order to assess baseline and laparotomy-induced effects on motor coordination, mice in the laparotomy groups and age-matched non-surgical controls were trained to stay on an accelerating rotarod during 6 pre-operative trials followed by evaluation of their rotarod performance after laparotomy on POD 2 and 4. During the training phase (pre-operative trials 1-6), an age-dependent difference in baseline accelerating rotarod performance was observed when comparing latency to fall times of 20- and 24-month-old mice to 3-month-old mice. (Figure 1C) Specifically, the latency to fall times of 24-month-old control mice were shorter in all pre-operative trials (p<0.02) and shorter in the 20-month-old control mice during trials 1-3 (p<0.004) but not in trials 4-6, compared to 3-month-old mice. These data suggest that while both the 20- and 24-month-old control mice were able to learn the rotarod task with training, their baseline performance was consistently poorer compared to 3-month-old mice. Despite this difference in baseline performance, the 20-month-old mice appeared to perform the rotarod task at a level approaching that of the 3-month-old mice after 3 training trials, suggesting that 20-month-old, but not 24-month-old mice have the physical capacity to perform the rotarod protocol tested similar to that of young mice.

Laparotomy did not reduce latency to fall times on POD 2 and 4 when comparing mice that underwent surgery to controls for each of the age groups tested. Latency to fall times trended toward a progressive decline on POD 2 and 4 in the 24-month-old mice after laparotomy but did not reach the threshold for statistical significance.

Age-dependent differences in baseline rotarod performance and training response were observed when comparing latency to fall times of 20-month-old (trials 1-4, p<0.02) and 24-month-old (trials 1-6, p<0.01) mice to 3-month-old mice in the laparotomy groups. In mice that underwent laparotomy, 24-month-old mice had a shorter latency to fall time on POD 2 and 4 (p<0.01) compared to 3-month-old mice, and on POD 4 (p=0.02) compared to 20-month-old mice. Taken together, these findings indicate that motor coordination in mice declines with increasing age in general and suggest a greater decrement in rotarod performance in response to laparotomy in 24-month-old mice.

### Differences in pre-operative motor coordination did not predict physical function measures after laparotomy in 24-month-old mice

Improvement in performance on the accelerating rotarod during pre-operative training trials 1-6 was highly variable in 24-month-old mice. Since baseline physical function in humans may affect surgical outcomes, the 24-month-old mice were subsequently designated as low functioning (LF) or high functioning (HF) based upon their performance on the rotarod during pre-operative trials 1-6. Mice with an average latency to fall time below the group mean were categorized as LF while mice with an average latency to fall time above the group mean were categorized as HF. Forelimb grip strength was then analyzed using the LF and HF group designations. The laparotomy procedure induced a reduction in average grip strength in both LF and HF groups (p<0.0001) similar to the aggregate result for all 24-month-old mice. (Figure 2A) Moreover, grip strength was similar between the LF and HF control mice, and similarly reduced between LF and HF mice that underwent laparotomy. These findings suggest that differences in baseline motor coordination in 24-month-old mice did not differentially affect post-laparotomy grip strength performance.

**Figure 2:**
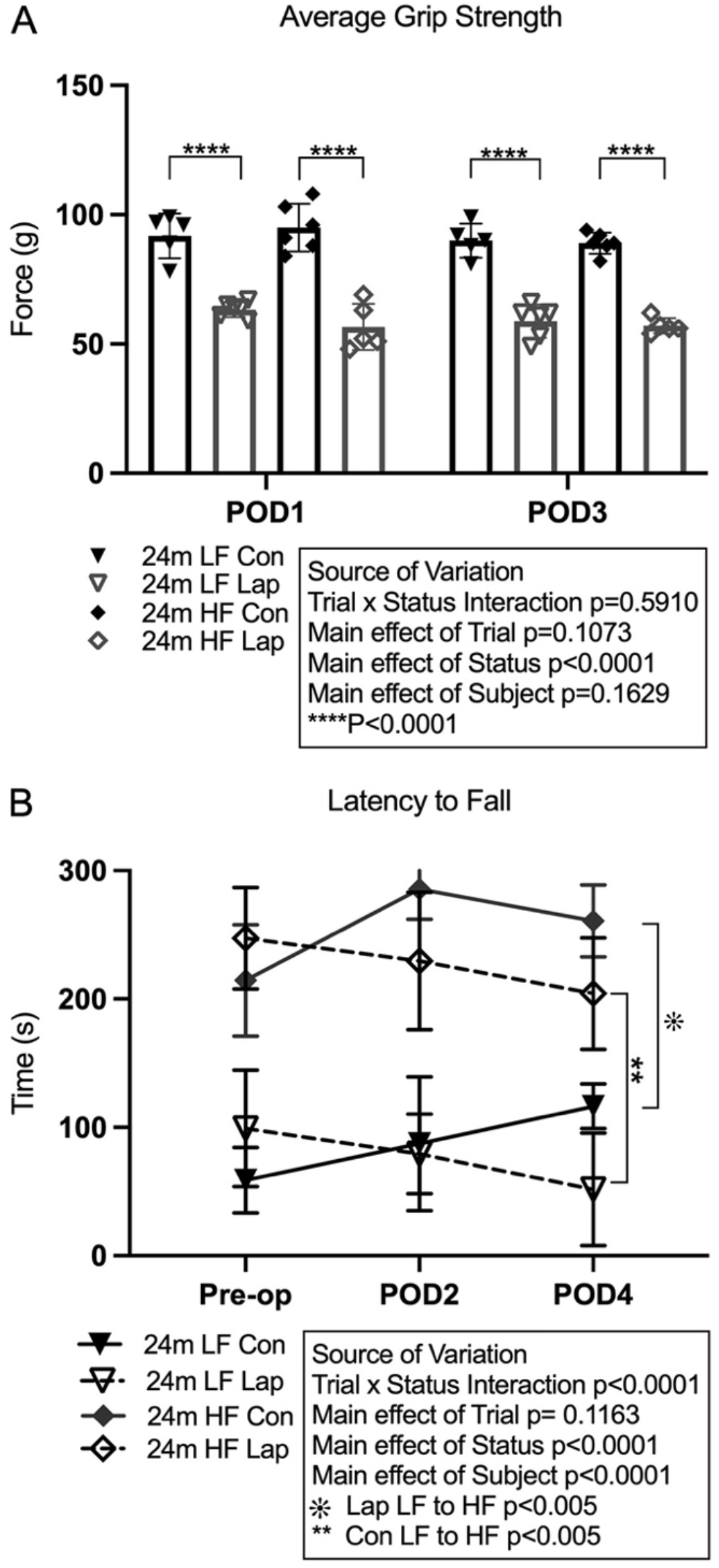
Behavioral analysis of 24-month-old C57BL/6N mice grouped by baseline performance on the accelerating rotarod. Low functioning (LF) and high functioning (HF) groups correspond to mice with an average latency to fall time below, or above, the group mean, respectively. (A) Comparison of the average grip strength force between the LF and HF control and laparotomy groups on POD 1 and 3. (B) Comparison of performance on the accelerating rotarod for LF and HF control and laparotomy groups during pre-operative trial and on POD 2 and 4. N=5-6 mice/group. Statistical analysis performed by 2-way ANOVA with Tukey test for multiple comparisons; a repeated measures design was used for rotarod analysis. Error bars denote standard deviation. POD=post-operative day; LF=low functioning; HF=high functioning.

The 24-month-old HF mice had longer latency to fall times compared to LF mice, in both the control and laparotomy groups, during the pre-operative trials (trials 1-6) and on POD 2 and 4 (p<0.005). (Figure 2B) However, laparotomy did not significantly reduce latency to fall times on POD 2 and 4 in either LF or HF groups. Thus, differences in baseline motor coordination in 24-month-old mice did not predict post-laparotomy rotarod performance.

### Laparotomy induced aged-associated reduction in bodyweight but not muscle mass

Laparotomy did not cause a reduction in body weight in 3-month-old mice during the post-operative period (POD 1-5). Following laparotomy, the 20-month-old and 24-month-old mice had a decrease in body weight, calculated as the percentage change in weight compared to pre-operative weight, on POD 1-3. (Figure 3A) The decline in body weight persisted on POD 4 but not on POD 5 in the 24-month-old laparotomy group. In the 20-month-old laparotomy group there was a decline in body weight on POD 5 but not on POD 4. Additionally, mice in the 20- and 24-month-old laparotomy groups had a greater loss in the percentage of body weight after surgery compared to mice in the 3-month-old laparotomy group (p=0.0352 and p=0.0185, respectively) on POD 2.

**Figure 3:**
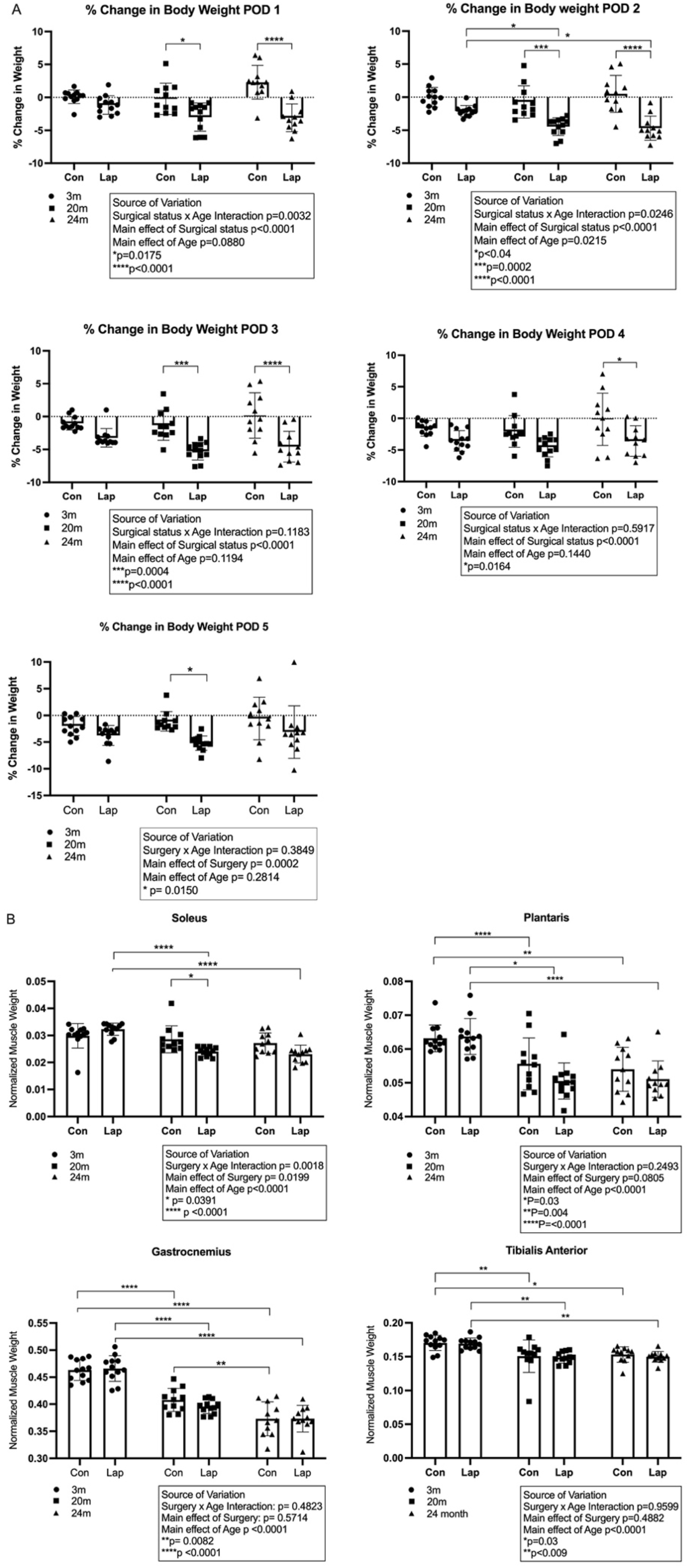
Laparotomy induced aged-associated reduction in bodyweight but not muscle mass in C57BL/6N mice. (A) Percentage change in body weight on POD 1-5 compared to pre-operative body weight. (B) Muscle weights of soleus, plantaris, gastrocnemius and tibialis anterior normalized to body weight on POD 5. N=11-12/group. Statistical analysis performed by 2-way ANOVA with Tukey test for multiple comparisons. Error bars denote standard deviation. POD=post-operative day; Con=control; Lap=laparotomy.

The weights of soleus, plantaris, gastrocnemius, and tibialis anterior muscles harvested on POD 5 were normalized to pre-operative body weights and compared between the groups of mice. (Figure 3B) In the control groups, 20- and 24-month-old mice had lower plantaris, gastrocnemius, and tibialis anterior muscle weights compared to 3-month-old mice. Furthermore, the 24-month-old mice had lower gastrocnemius muscle weight compared to 20-month-old mice (p=0.0082). In the mice that underwent laparotomy, 20- and 24-month-old mice had lower soleus, plantaris, gastrocnemius, and tibialis anterior muscle weights compared to 3-month-old mice. Of these muscles, only the soleus muscle of the 20-month-old mice showed a reduction in muscle mass after laparotomy compared to age-matched controls (p=0.039).

Quantification of food intake showed decreased food consumption in the 20-month-old laparotomy group compared to non-surgical controls on POD 1 (p=0.0049) but not on POD 2-5. (Supplemental Figure 1) Laparotomy did not induce a reduction in food intake post-operatively (POD 1-5) compared to control mice in either 3 or 24-month-old mice.

### Laparotomy did not affect protein synthesis in soleus and plantaris muscles

Global protein synthesis quantification using the SUnSET technique was performed on soleus and plantaris muscles collected on POD 5 from 3-, 20- and 24-month-old mice following laparotomy and their age-matched controls. There were no Surgery x Age interaction effects for global protein synthesis observed for the soleus (p=0.179) or plantaris (p=0.691). We did note a main effect of age in the soleus muscles (p=0.012), with the 20-month-old mice having elevated protein synthesis compared to the 3-month-old mice (p=0.011) and a modest trend for greater protein synthesis in the 24-month-old mice compared to 3-month-old mice (p=0.077). These findings suggest that laparotomy does not induce significant changes in protein synthesis in soleus and plantaris muscles 5 days after surgery. (Supplemental Figure 2)

### Laparotomy induced age-associated differential gene expression and canonical pathway activation in skeletal muscles

RNA sequencing (RNA-seq) studies were performed using total RNA isolated from soleus muscles collected on POD 5. Differential gene expression was determined by comparing mice that underwent laparotomy to their respective age-matched controls. Laparotomy had a greater effect on the number of differentially expressed genes (DEGs) in 3-month-old C57BL/6N mice (997 DEGs) compared to 20-month-old (139 DEGs) and 24-month-old mice (75 DEGs). (Figure 4A) When the 24-month-old mice were compared based on their baseline performance on the rotarod, the LF and HF groups had similar numbers of DEGs (228 and 224, respectively), and yet only 21 DEGs were common between the two comparisons. (Figure 4B) The common genes with similar expression patterns, including *Pparα, Fst, Myh4*, and *Serpine1*, may be hallmarks of the post-surgical molecular response associated with aging. In contrast, the common genes with reversed expression patterns (*Adamtsl1, Comp, Nmrk2, Pla2g15* and *Tmsb10*) may provide insight into the key factors that separate the LF and HF groups. See Supplemental Files for gene lists of DEGs in soleus muscles of 3-month, 20-month, 24-month, 24-month LF and 24-month HF mice.

**Figure 4:**
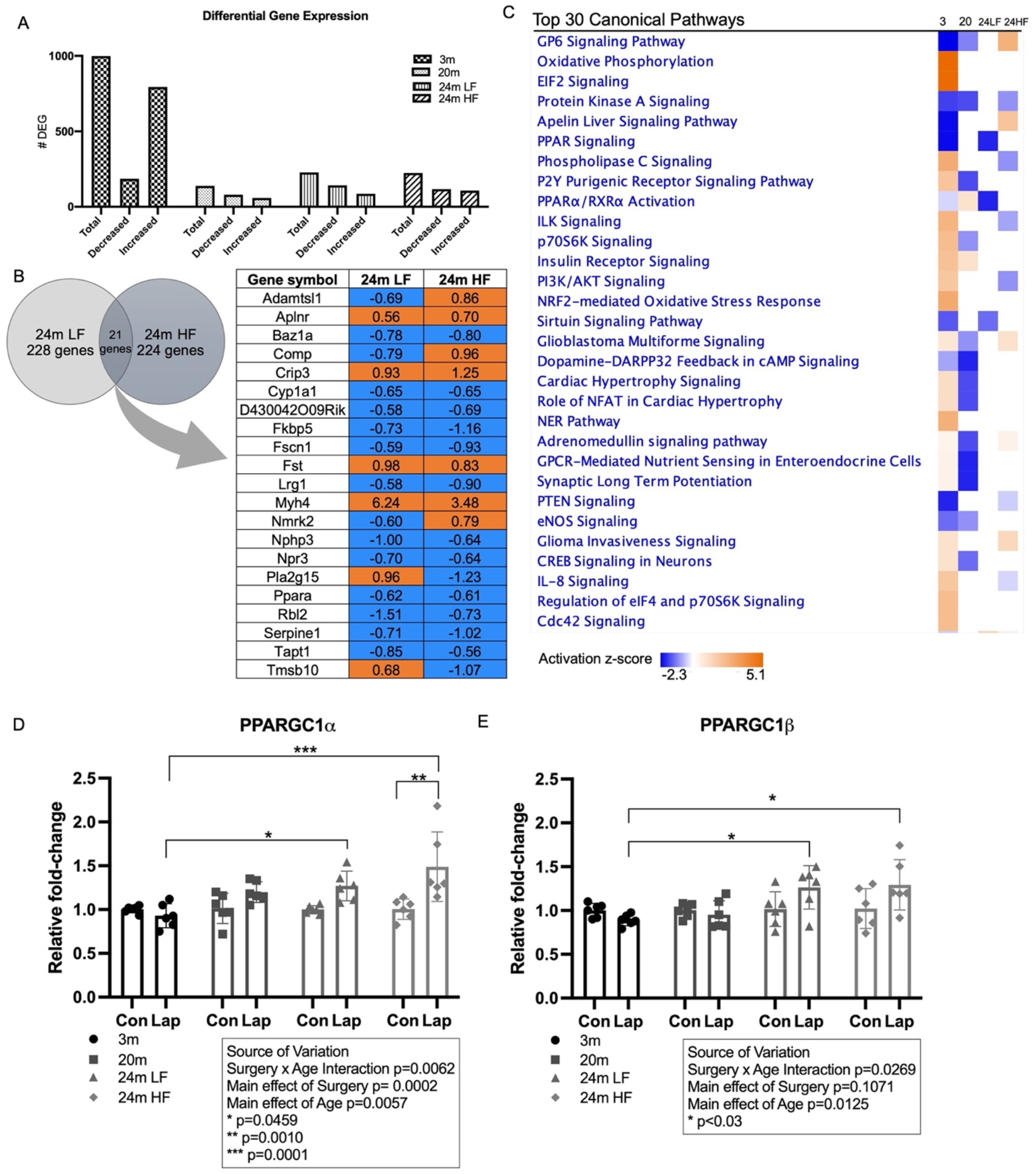
RNA-seq and pathway analysis show an age-associated effect on skeletal muscle gene expression after laparotomy. (A) The number of DEGs in soleus muscles harvested 5 days following laparotomy vs age-matched controls, N=3/group. (B) DEGs that are common between 24-month-old LF and HF groups. (C) Heatmap of the top 30 canonical pathways affected by laparotomy ranked by z-score. (D and E) Expression of *PPARGC1α* and *PPARGC1β* in the tibialis anterior muscle as measured by RT-PCR, N=6/group. Statistical analysis performed by 2-way ANOVA with Tukey test for multiple comparisons. Error bars denote standard deviation. Con=control; Lap=laparotomy; LF=low functioning; HF=high functioning; DEG=differentially expressed gene.

Pathway analysis was performed using Ingenuity Pathway Analysis (IPA) to understand the functional implications of these DEGs in 3-, 20-, and 24-month-old LF or HF mice. The analysis showed that, as compared to 20- or 24-month-old mice, a greater number of canonical pathways were affected in 3-month-old mice following laparotomy. (Figure 4C) The comparison analysis showed robust activation of oxidative phosphorylation (*Atp5e, Ndufa13, Ndufa2, Uqcrq*), phospholipase C signaling (*Hras, Map2k2, Prkcz, Myl3*), and NRF2-mediated oxidative stress response (*Sod1, Sod3, Hras, JunD*) pathways in the 3-month-old mice following laparotomy, but not in the older mice. Other pathways in the 3-month-old mice were inhibited including PPAR signaling (*Hras, Map2k2, Jun, Nfkbib*), PPARα/RXRα activation (*Hras, Map2k2, Jun, Jak2*), and PTEN signaling (*Hras, Map2k2, Prkcz, Csnk2b*). Inhibition of several of these pathways was also noted in the 24-month-old LF mice but DEGs responsible for pathway inhibition were different [e.g., PPAR signaling (*Fos, Ppargc1a, Pparg, Ppara*) and PPARα/RXRα activation (*Ppara, Ppargc1a, Adipoq, Slc27a1*)].

Upstream regulator analysis was performed using IPA to identify candidate transcriptional regulators that might explain the observed gene expression changes identified via RNA-seq. Several putative upstream regulators were revealed. To explore if any of these candidates might participate in the gene expression changes, mRNA expression of these regulatory factors was quantified by RT-PCR. Of the 15 upstream targets evaluated (Supplemental Table 1), the expression of *PPARGC1α* and *PPARGC1β* were significantly increased in tibialis anterior muscles of the 24-month-old LF and HF mice that underwent laparotomy compared to the 3-month-old laparotomy group. (Figure 4D and 4E) Additionally, laparotomy induced increased expression of *PPARGC1α* in tibialis anterior muscles of the 24-month-old HF mice that underwent laparotomy compared to age-matched controls.

### Lipidomics analysis of serum and gastrocnemius muscle

Because post-operative changes in LMs were anticipated,[15] we used a liquid chromatography-mass spectrometry (LC-MS) based lipidomics approach to profile changes in serum and muscle levels of LMs after laparotomy to examine the effects of age and function on such changes. Altered lipidomic profiles of serum and gastrocnemius muscle were observed from 3-month-old and 24-month-old LF and HF mice. (Figure 5). Serum levels of lipid mediators (LMs) derived from ω-3 polyunsaturated fatty acids (PUFAs), such as eicosapentaenoic acid (EPA), docosahexaenoic acid (DHA) and α-linolenic acid (ALA), were, in general, reduced slightly by aging (3-month-old vs. 24-month-old LF and HF mice), while metabolites from ω-6 PUFAs like arachidonic acid (AA) increased in mouse serum with age (Figure 5A). Unlike other hydroxyeicosapentaenoic acids (HEPEs) and hydroxyeicosatetraenoic acids (HETEs), 12-HEPE and 12-HETE, generated through 12-lipoxygenase (12-LOX) from EPA and AA respectively, exhibited markedly higher serum levels in 24-month-old LF/HF groups than 3-month-old mice. LM levels in gastrocnemius were associated with the rotarod performance on pre-operative trials 1-6 (i.e., baseline functioning). When compared with 3-month-old control mice, LMs synthesized through EPA, DHA, ALA, linoleic acid (LA) and AA/LOX pathways decreased in 24-month-old control mice from the LF group, whereas these LMs increased in 24-month-old control mice from the HF group (Figure 5B).

**Figure 5:**
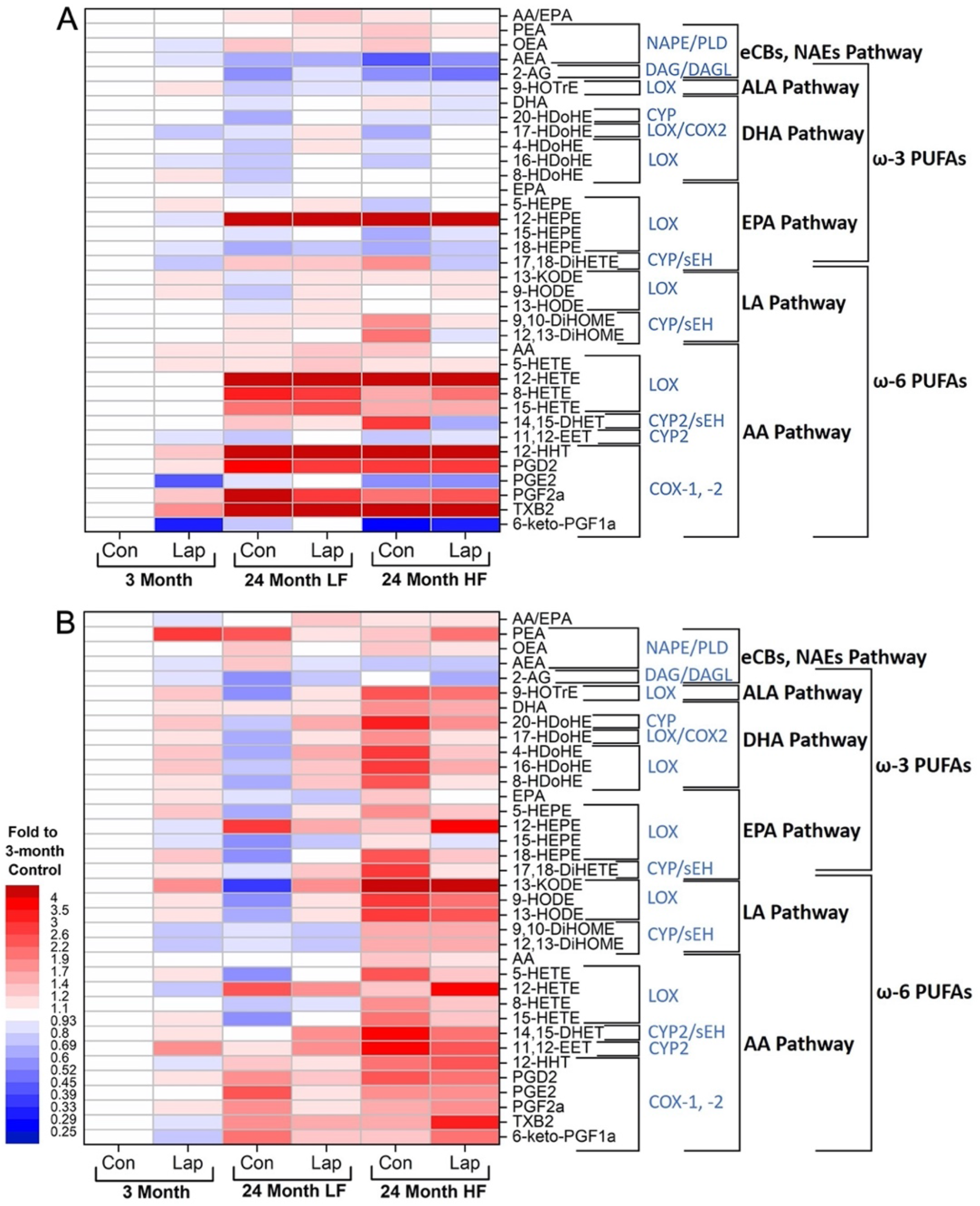
Heatmaps of lipidomic profiles in mouse serum and gastrocnemius muscle suggest that age and function may change the levels of lipid signaling mediators derived from different metabolic pathways. (A) Heatmap of lipid mediators in serum. (B) Heatmap of lipid mediators in gastrocnemius muscles. N=3/group; Con=control; Lap=laparotomy; LF=low functioning; HF=high functioning.

Further statistical analysis using 2-way ANOVA was performed to reveal differences in the main effects of four lipid mediators in both sets of samples (Supplemental Figure 3A and 3B). The concentration of thromboxane (TXB_2_) and 12-hydroxyheptadecatrienoic acid (12-HHT), two lipid mediators derived from AA through the cyclooxygenase (COX) pathway, showed a difference in the main effect of age/functional status in both serum (TXB_2_ p=0.0125 and 12-HHT p=0.0111) and gastrocnemius muscle (TXB_2_ p=0.0024 and 12-HHT p=0.0273). In the gastrocnemius muscle of mice that underwent laparotomy, TXB_2_ concentration was higher in the 24-month-old HF mice compared to 24-month-old LF mice (p<0.044) and the 3-month-old mice (p=0.0026). Additionally, there was an increase in TXB_2_ concentration in the gastrocnemius muscle of the 24-month-old HF mice after laparotomy compared to age-matched controls (p=0.044).

Anandamide (AEA) and palmitoylethanolamide (PEA), two signaling lipids belonging to endocannabinoids/N-acylethanolamines (eCBs/NAEs) family, showed a difference in the main effect of age/functional status in serum (p=0.0434 and p=0.0227, respectively) when analyzed using 2-way ANOVA. The concentration of PEA in the gastrocnemius muscle showed a difference in the main interaction between the variables of surgery and age (p=0.0113). The AEA concentration in the gastrocnemius muscle was lower in 24-month-old HF control mice compared to 24-month-old LF control mice (p=0.0209). Otherwise, no significant differences in AEA or PEA concentrations were observed between any other groups.

## Discussion

In older adults, age and pre-operative physical functioning are predictors of adverse post-operative outcomes.[16, 17] However, the biological mechanisms underlying late-life vulnerability to post-operative physical decline remain poorly understood. In this study, we characterized the effects of a laparotomy procedure on measures of post-operative physical function and skeletal muscle gene expression in C57BL/6N mice across life span. We found that laparotomy induced age-associated decline in muscle strength and differential regulation of expression of mRNAs encoding proteins participating in canonical pathways in skeletal muscles after surgery. In addition, lipidomics studies in gastrocnemius muscles confirmed the associations between profiles of signaling lipids and age/functional status in mice.

Our laparotomy model showed an age-associated post-operative decline in forelimb grip strength in 3, 20 or 24-month-old mice with the oldest mice showing the greatest reduction and the slowest recovery in grip strength after surgery. These findings are consistent with the patterns of post-operative handgrip strength decline and recovery after abdominal surgery in older adults in whom the rate of muscle strength recovery is less rapid and complete compared to younger adults.[7] The cause of post-operative decline in forelimb muscle strength in our mouse model is unclear. However, it does not appear to be linked to a reduction in skeletal muscle protein synthesis in the lower limb muscles studied (as indicated by SUnSET) or loss of muscle mass via catabolism (as inferred from the general lack of differences in skeletal muscle weights in the muscles studied and similar food intake). Previously, Edwards and colleagues observed that muscle endurance and isometric strength were reduced post-operatively in a small group of human subjects.[6] Thus, studies utilizing ex-vivo muscle physiology technique are needed to evaluate whether laparotomy exacerbates age-associated slowing of muscle contraction speed and diminishes muscle power production, both of which contribute to muscle dysfunction in the absence of loss of muscle mass in aged animals [18, 19]. Interestingly, despite pre-operative differences in motor coordination between the 24-month-old LF and HF mice, baseline functional status did not affect the severity of decline in laparotomy-induced forelimb grip strength in these mice. It is possible that gross motor skills such as movement and coordination, while impacting large muscle groups and whole-body movement, do not have a substantial effect on muscle strength in the forelimbs of aged mice.

While age-associated differences in baseline motor coordination and ability to acquire rotarod task were observed in mice 3, 20 or 24 months of age consistent with previously published studies,[20, 21], laparotomy did not significantly impair post-operative rotarod performance in these mice. However, a trend towards poorer post-operative motor coordination up to 4 days after laparotomy was noted in the 24-month-old mice. Moreover, it appears that the laparotomy procedure modeled in our experiments and behavioral testing paradigm negatively affects post-operative skeletal muscle strength but not motor coordination and that baseline motor coordination in aged mice is unrelated to post-operative skeletal muscle strength. These observations likely reflect the complexity of motor coordination in which an intricate set of physiological controls of movement with brain, spinal cord, nerve and neuromuscular junctions, as well as sensory structures for joint position and muscle and tendon loading fine tune the neural output. Thus, surgery did not seem to alter these higher-order determinants of muscle function in this mouse model although further study is required to clearly prove this conclusion. Taken together, these findings point to the skeletal muscle as the vulnerable target organ in this model system.

RNA sequencing studies showed robust age-associated effects on post-operative skeletal muscle gene expression. At 5 days post laparotomy, the soleus muscles of 3-month-old mice had a substantial number of DEGs induced by the surgical stress from laparotomy. This profound skeletal muscle response to the surgical insult may represent an attempt to preserve muscle function and reestablish homeostasis following a traumatic insult in young mice. The sizeable differences in the number of DEGs observed between the 3-month-old and the older (20- and 24-month-old) mice may be indicative of a maladaptive post-operative tissue response to surgical stress in the skeletal muscles of the older mice 5 days after laparotomy. It is possible that the low number of DEGs in response to laparotomy in older mice reflects impaired homeostatic mechanisms operating through altered gene expression that is either unresponsive to, or is already maximally stimulated by, the aged organism’s response to metabolic stress after surgery. Examples of such aging-associated, differentially regulated signaling pathways that were activated in the 3-month-old, but not older mice include canonical pathways for oxidative phosphorylation, phospholipase C signaling, and NRF2-mediated oxidative stress response. Interestingly, while the 24-month-old LF and HF mice had similar total numbers of DEGs after laparotomy, only 21 of these genes were common between the two groups. It may be possible that some of these genes (e.g., *Pparα, Fst, Myh4* and *Serpine1*) constitute hallmarks of post-surgical molecular response in skeletal muscles of aged organisms. It is noteworthy that these genes respectively represent fundamental cellular signaling pathways driving metabolic adaptations, telomerase function, muscle contraction, and cell adhesion/ECM remodeling. To our knowledge, this study is the first to explore the relationship between age, physical function, and patterns of skeletal muscle gene expression after surgical stress. While the findings are descriptive, they inform future investigations aimed to determine molecular mechanisms that drive surgery-induced muscle and functional decline in aged organisms.

Analysis of levels of lipid signaling molecules in skeletal muscles (gastrocnemius) also confirmed the associations between such profiles and age/functional status. Lipid mediators can regulate skeletal muscle mass and function and potentially lead to muscle wasting in response to various physiological and pathological conditions.[22, 23] It is believed that ω-3 and ω-6 PUFAs have opposing effects on metabolic functions in the body. ω-6 PUFAs such as AA and its metabolites prostaglandins (PGs), thromboxanes (TXs), and others are generally associated with inflammatory responses, constriction of blood vessels, and platelet aggregation.[24-26] In contrast, ω-3 PUFAs including EPA, DHA and related metabolites, show anti-inflammatory and pro-resolving activities.[27, 28] In this study, 24-month-old mice with LF showed reduced levels of lipid mediators related with ω-3 PUFAs (EPA, DHA, and ALA) but markedly higher levels of PGs and TXs, a group of ω-6 AA metabolites generated via COX metabolic pathways, as compared with young mice, suggesting a age-related increased role for lipid mediators in inflammation in skeletal muscles during aging. However, higher ω-3 and lower ω-6 PUFAs (PGs, TXs) levels were observed in the aged mice (24-month-old) with higher functioning than those with lower functioning, suggesting that physical function status may potentially affect inflammatory response at the same age. Yet, another intriguing possibility is that low and high functioning mice might respond differently at the epigenetic level to lipid mediator signaling.[29]

This study has several limitations. The C57Bl/6N mice, particularly the 24-month-old animals, had a wide range of rotarod performance abilities suggesting differences in motor coordination skills and task acquisition. We attempted to mitigate this variability by dividing the 24-month-old mice into low and high functioning groups based upon their pre-operative rotarod performance abilities. While this approach differentiated baseline rotarod performances between LF and HF mice and helped to identify gene expression differences post-operatively, it reduced the group size and power to detect changes in post-operative physical performance outcomes. Moreover, although balance and coordination as measured by accelerating rotarod assessed one important aspect of physical function that declines with aging, a more comprehensive approach to functional assessment may be necessary to better delineate pre-operative functional status of the 24-month-old mice [30]. Lastly, the inclusion of behavioral tests that pose a greater post-operative physiological challenge such as forced treadmill running will be needed in future studies to better capture differences in physical performance abilities in LF and HF mice.

In summary, findings from this study provided evidence that a mouse model of laparotomy recapitulates some features of post-operative skeletal muscle decline observed in older adults who undergo surgery, and identified age-associated, laparotomy-induced molecular signatures in skeletal muscles. Thus, future studies can build upon this approach to further investigate molecular mechanisms of late-life vulnerability to acute surgical stress and resiliency to surgery-induced physical decline.

## Methods

### Animals

C57BL/6N mice from the NIA Aged Rodent Colonies [31] were individually housed in the Veterinary Medical Unit at the James J. Peters VA Medical Center (JJPVAMC) in rooms that provided a controlled temperature of ~72º F and a 12:12 hour light:dark cycle. Animals were allowed to acclimate to their home cages and were given food and water *ad libitum* prior to the beginning of the study. The study protocol was approved by the JJPVAMC Institutional Animal Care and Use Committee.

### Food Intake

At the outset of the study, the food hoppers were removed from the mouse cages. Each mouse was given 10-13 grams of rodent chow daily placed on the bottom of the cage. The weight of the chow was recorded each morning. The following morning, the remaining chow was removed and weighed to determine the amount of food consumed in the previous day.

### Body Weights

All mice were weighed the morning of the laparotomy and daily during the post-operative period to monitor for changes in body weight.

### Laparotomy surgery

Mice were anesthetized by inhalation of isoflurane (5% for induction and 3% for maintenance) for 15 minutes (3 minutes induction; 12 minutes maintenance via nose cone). After induction of anesthesia, the abdomen was shaved then swabbed with betadine solution. A 1 cm midline vertical mark was placed on the mid-abdomen to indicate the surgical site. Using a fine surgical scalpel (#11 blade), a 1 cm vertical (perpendicular to the skin) midline incision traversing the skin, subcutaneous tissue, muscles, and peritoneum was made. Then, the abdominal wall was closed with 4 interrupted 4-0 coated vicryl sutures. The skin and subcutaneous tissue were closed using the same technique. The laparotomy procedure lasted ~8 minutes. Body temperature was maintained at 37°C by laying the animal on a circulating water heating pad during the surgical procedure. Mice were given subcutaneous enrofloxacin (5 mg/kg of body weight) and carprofen (5 mg/kg of body weight) immediately prior to surgery. Mice were returned to their home cages following surgery and monitored until they were ambulatory. Carprofen (5 mg/kg of body weight) was administered on POD 1-3 for analgesia. Control mice received the same dose and course of treatment of enrofloxacin and carprofen as the laparotomy groups.

### Grip strength

Forelimb muscle strength was measured using a grip strength meter (Columbus Instruments, Columbus, OH) with a sensor range of 0–500 grams and accuracy of 0.15%.[32] Each mouse was held by the tail and allowed to grasp the metal grid of the apparatus with its front paws while steady pressure was applied to the tail. The grip strength measurement was taken just before release to determine maximum tensile force in grams. Three measurements were taken and averaged for each day of assessment (pre-operative trial and POD 1 and 3).

### Accelerating rotarod

Balance and motor coordination were assessed by an accelerating rotarod apparatus (IITC Life Science Inc, Woodland Hills, CA).[20] The mouse was placed horizontally on the rotating rod prior to the start of each session. Acclimation training was performed at a constant rod speed of 5 rpm for 2 minutes. On all subsequent days (pre-operative trial days 1 to 6, and POD 2 and 4), the rod rotation speed was programed to increase from 5 to 10 rpm over 5 minutes. The time the mouse was able to stay on the rotating rod before falling (i.e., latency to fall time) was determined up to a maximum duration of 300 seconds. Each mouse performed three sessions (one hour apart) per day, three days per week for the first two weeks (acclimation, pre-operative trial days 1 to 5), then two days during the third week (pre-operative day 6 and POD 2), and one day during the fourth and final week (POD 4; Figure 1A). The latency to fall time was recorded as the seconds elapsed between the start of the session and the time at which the mouse fell from the rotating rod. The average latency to fall time was calculated for each trial day and used to assess motor coordination performance. Control mice were subjected to the same rotarod protocol.

### Tissue collection

Mice were anesthetized with isoflurane (5% for induction, 3% for maintenance) on POD 5. Entire skeletal muscles (soleus, plantaris, gastrocnemius and tibialis anterior) were carefully dissected under anesthesia, and immediately weighed and flash frozen in liquid nitrogen chilled isopentane. Blood was collected via cardiac puncture then centrifuged at 12,000 g for 5 minutes. The serum was collected and frozen at -80°C.

### Biochemistry

See Supplemental Materials for detailed descriptions of biochemistry techniques used in this study.

### Surface sensing of translation (SUnSET)

On POD 5, the right soleus and plantaris muscles were carefully excised and placed in an organ bath solution of Krebs-Hanseleit, 1× MEM and 25 mM glucose for 15 minutes at 37°C, then rinsed in PBS and placed in serum-free DMEM with 10 µM puromycin (Calbiochem) for 30 minutes at 37°C.[33] Muscles were then snap frozen in liquid nitrogen for subsequent Western blotting. Frozen muscle was homogenized and centrifuged to collect the supernatant. 50 µg of protein was loaded into precast 4-15% gradient gels (Bio-Rad) and run at 180 V for 60 minutes. Proteins were transferred from the gel to a PVDF membrane using a semidry transfer and probed with a mouse monoclonal anti-puromycin primary antibody (Millipore, Cat# MABE343), followed by incubation with an anti-mouse secondary antibody (Jackson Immuno Research, Cat# 115-035-206). Membranes were incubated in a horseradish peroxidase chemiluminescent solution (GE Amersham), imaged using a CCD digital imager (Bio-Rad), and densitometry values of the complete lane were determined using Image Lab software (Bio-Rad) and normalized to whole lane densitometry values of the Ponceau S-stained membranes.

### RNA preparation

Frozen soleus or tibialis anterior muscle was homogenized using the FastPrep-24 5G benchtop homogenizer (MP bio). RNA was then prepared from the homogenate using the RNeasy Mini Kit (Qiagen).

### RNA sequencing and pathway analysis

Whole transcriptome analysis using RNA-seq was performed at the Genomics Core Lab, New York Medical College. Libraries were prepared with RNA extracted from soleus muscles using TruSeq Stranded Total RNA Library Prep Kit (Illumina). Sequencing of paired-end reads (75 bp × 2) was performed using the NextSeq 550 system (Illumina). Raw sequence reads were de-multiplexed and trimmed for adapters by using Illumina bcl2fastq conversion software (v2.19). Sequence reads of each sample was pseudo-aligned to Ensembl GRCm38 mouse reference transcriptome and the gene transcript abundance was quantified by using the Kallisto algorithm.[34] Differential gene expression was analyzed in paired groups by using DESeq2 package in RStudio (v1.1.423 with R v3.4.3).[35] The resulting data was filtered manually in Excel using the inclusion criteria of p ≤ 0.05, |log_2_(fold change) | ≥ 0.55, and median expression level ≥50. The filtered gene lists were uploaded to IPA (QIAGEN Inc., https://www.qiagenbioinformatics.com/products/ingenuity-pathway-analysis), in which pathway analyses, gene ontology and upstream regulator analyses were performed. All the RNA-seq data have been submitted to Gene Expression Omnibus (accession number GSE184486).

### cDNA synthesis and real time-PCR

cDNA was prepared from soleus and tibialis anterior muscle RNA using the High-Capacity cDNA reverse transcription kit (Applied Biosystems). Real time-PCR was performed using ViiA7 Real-Time PCR system (Applied Biosystems) with TaqMan gene expression assays (Applied Biosystems) using standard techniques. All gene expression assays were normalized to α-tubulin (*Tuba4a*). Relative gene expression levels were calculated using the -2^ΔΔCt^ method [36]. (See Supplemental Table 1 for a list of TaqMan gene expression assays used)

### Lipidomics

#### Tissue Preparation

50-100 mg of frozen gastrocnemius muscle was minced and homogenized using a Tissue Lyser II homogenizer (Qiagen). The muscle homogenate was mixed with 5 µl of isotope-labelled LM internal standards (IS) stock solution followed by centrifugation to remove tissue residue and precipitated proteins. An aliquot (50 µl) of serum from each mouse was mixed with 1.0 ml of ice-cold 80% methanol in water (v/v) containing 5 µl of isotope-labelled LM IS stock solutions followed by centrifugation to remove the precipitated proteins. All tissue samples were cleaned and concentrated by Solid Phase Extraction (SPE) before being injected into LC-MS/MS. The LMs from the SPE sorbent bed were eluted by methanol.

#### Sample Profiling

The LC separation was conducted on a C8 column (Ultra C8, 150 × 2.1mm, 3 µm, RESTEK) and the MS/MS analysis was performed on Shimadzu LCMS-8050 triple quadrupole mass spectrometer. We tested for levels of 158 lipid mediators (Shimadzu Scientific Instruments, Inc.,) and further optimized our previously published quantification method [37-39], including the m/z transitions (parents to product ions) and their tuning voltages based on the best MRM responses from instrumental method optimization software. All analyses and data processing were completed on Shimadzu LabSolutions V5.91 software.

### Statistical Analysis

All data are presented as means ± standard deviation. Muscle weights and body weights were normalized to the pre-operative body weights of mice prior to analysis. Two-way analysis of variance (ANOVA) with Tukey test for multiple comparisons was performed for all analyses except the rotarod for which a 2-way repeated measure ANOVA was used. Statistical analysis was performed using GraphPad Prism (version 9).

## Supporting information

Supplemental File 3 month old gene list

Supplemental File 20 month old gene list

Supplemental File 24 month old HF gene list

Supplemental File 24 month old LF gene list

Supplemental File 24 month old gene list

## Disclaimer

The work reported herein does not represent the views of the US Department of Veterans Affairs or the US Government.

## Acknowledgements

The authors wish to acknowledge Dr. Greg Elder for many helpful discussions during planning of these studies and data interpretation and the generous support of the James J Peters VA Medical Center. Author Contributions: Study Design: FK, CC, CM, ZG, SA and MB; Data acquisition: FK, ZG, SA, WH and ZW; Animal surgery and tissue/blood collection: FK, ZG and LH; Statistical analysis: FK, ZG, SA, AI and ZW; Data analysis and interpretation: FK, CC, CM, ZG, SA, MB, AI, WH and ZW; Manuscript writing: FK, CC, ZG, SA, MB and WH.

## Data Availability Statement

The authors declare that the data supporting the findings in this study are included in the paper and the supplementary information are available upon request from the corresponding author.

