## Supplemental Methods for "RNA-sequencing reveals a gene expression signature in skeletal muscle of a mouse model of age-associated post-operative functional decline"

**Surface sensing of translation (SUnSET):** On POD 5, the right soleus and plantaris muscles were carefully excised, weighed, and placed in ice cold PBS. The muscles were then placed in an organ bath solution of Krebs-Hanseleit, 1× MEM and 25 mM glucose for 15 minutes at 37˚C with moderate shaking at 100 rpm on an orbital shaker. The muscles were rinsed in PBS and then placed in serum-free DMEM with 10 µM puromycin (Calbiochem) for 30 minutes at 37˚C with moderate shaking at 100 rpm. Muscles were then snap frozen in liquid nitrogen.[33]

Frozen soleus or plantaris muscles was homogenized using a Dounce homogenizer in a cocktail of 1× RIPA buffer with protease and phosphatase inhibitors. Samples were incubated on ice for 30 minutes with 10 seconds of vigorous vortexing every 10 minutes. The homogenate was spun down at 4˚C for 15 minutes at 14,000 g, and the supernatant was collected. Protein concentration was determined using a microBCA kit (Pierce). 50 µg of protein was loaded into precast 4-15% gradient gels (Bio-Rad) and ran at 180 V for 60 minutes. Proteins were transferred from the gel to a PVDF membrane using a semidry transfer at 20 V for 50 minutes. Membranes were stained with Ponceau S, imaged using a CCD digital imager, then de-stained for 10 minutes in Tris-Buffer Saline with 0.1% Tween 20 (TBST). Membranes were blocked with 5% BSA for one hour at room temperature and then incubated overnight in a 1% BSA-TBST solution containing a mouse monoclonal anti-puromycin primary antibody (Millipore, Cat# MABE343). After overnight incubation, membranes were washed for 3 × 10 minutes in TBST and placed in 1% milk-TBST with an anti-mouse secondary antibody (Jackson Immuno Research, Cat# 115-035-206) for 60 minutes at room temperature. Membranes were then washed for 3 × 10 min in TBST, incubated for 5 minutes in a horseradish peroxidase chemiluminescent solution (GE Amersham), and imaged using a CCD digital imager (Bio-Rad). Densitometry values of the complete lane were determined using Image Lab software (Bio-Rad) and were normalized to whole lane densitometry values of the Ponceau S-stained membranes. To limit blot-to-blot variability, each sample was further normalized to a control sample of tibialis anterior muscle exposed to the SUnSET protocol but not part of the current study which was also run on each gel.

**RNA preparation:** Frozen soleus or tibialis anterior muscle was placed into a 2 mL lysing matrix tube containing a ¼” ceramic bead (MP bio) with 700 µL of QIAzol lysis reagent (Qiagen). The muscle was homogenized for 40 seconds using the preprogramed mouse muscle setting on the FastPrep-24 5G benchtop homogenizer (MP bio). RNA was then prepared from the muscle homogenate using the RNeasy Mini Kit (Qiagen). An on-column DNase digestion was performed using the RNase-Free DNase set (Qiagen) to ensure removal of DNA from samples. RNA concentration was measured with a NanoDrop 1000 spectrophotometer (Thermo Scientific). Samples were considered to be free from contamination when the 260/280 nm ratio was ~2.0.

**RNA sequencing:** Whole transcriptome analysis using RNA-seq was performed at the Genomics Core Lab, New York Medical College. Libraries were prepared with RNA extracted from soleus muscles using TruSeq Stranded Total RNA Library Prep Kit (Illumina). Sequencing of paired-end reads (75 bp × 2) was performed using the NextSeq 550 system (Illumina). Raw sequence reads were de-multiplexed and trimmed for adapters by using Illumina bcl2fastq conversion software (v2.19). Sequence reads of each sample was pseudo-aligned to Ensembl GRCm38 mouse reference transcriptome and the gene transcript abundance was quantified by using the Kallisto algorithm.[34] Differential gene expression was analyzed in paired groups by using DESeq2 package in RStudio (v1.1.423 with R v3.4.3).[35]

**Gene expression and pathway analysis:** The data from the RNA-seq analysis was filtered manually in Excel using the inclusion criteria of p ≤ 0.05, |log_2_(fold change) | ≥ 0.55, and median expression level ≥50. The filtered gene lists were uploaded to IPA (QIAGEN Inc., https://www.qiagenbioinformatics.com/products/ingenuity-pathway-analysis), in which pathway analyses, gene ontology and upstream regulator analyses were performed. All the RNA-seq data have been submitted to Gene Expression Omnibus with accession number GSE184486.

**cDNA synthesis and real time-PCR:** cDNA was prepared from soleus and tibialis anterior muscle RNA using the High-Capacity cDNA reverse transcription kit with RNase inhibitor (Applied Biosystems) according to the manufacturer’s instructions. One microgram of RNA was used for each cDNA reaction of 20 µL. cDNA samples were diluted with 80 µL of RNase-free water then stored at -20˚C.

Real time-PCR was performed using TaqMan gene expression assays (Applied Biosystems). Each reaction of 20 µL was prepared containing 1 µL of primers and probe mixture, 10 µL of 2× TaqMan universal master mix II, no UNG (Applied Biosystems), 2 µL of cDNA template, and 7 µL of RNase-free water. The reactions were carried out in a 96-well plate with the ViiA7 Real-Time PCR system (Applied Biosystems), programed as per the manufacturer’s instructions for the TaqMan gene expression assays. All gene expression assays were normalized to α-tubulin (*Tuba4a*) reference and the relative gene expression levels were calculated using the -2^ΔΔCt^ method [36]. Please refer to Supplemental Table 1 for a list of TaqMan gene expression assays used in this study.

**Lipidomics:**

**Tissue Preparation:** 50-100 mg of frozen gastrocnemius muscle stored at -80°C was defrosted on ice and minced into small pieces. The minced muscle and one stainless steel bead (5 mm, Qiagen) were placed into a 2 ml round-bottom low retention microcentrifuge tube (Fisher Scientific) then 1 ml of ice-cold 80% methanol in water (v/v) was added. The mixture was homogenized using a Tissue Lyser II homogenizer (Qiagen) at the frequency of 30 s^-1^, in 8×30-s bursts, waiting 20 s in between to avoid heating the samples. The muscle homogenate was mixed with 5 µl of isotope-labelled LM internal standards (IS) mixture stock solution (5 µg/ml for AA-d_8_, 2 µg/ml for DHA-d_5_ and EPA-d_5_, and 0.5 µg/ml for the rest IS), and then agitated on ice in the dark for 1-2 hours, followed by centrifugation at 16,000×g at 4°C for 10 minutes to remove any tissue residue and precipitated proteins.

Aliquot (50 µl) of serum from each mouse was mixed with 1.0 ml of ice-cold 80% methanol in water (v/v) containing 5 µl of isotope-labelled LM IS stock solutions which was same as gastrocnemius muscle preparation, and then agitated on ice and in the dark for 15 min, followed by centrifugation at 16,000×g at 4°C for 10 min to remove the precipitated proteins.

All tissue samples were cleaned and concentrated by Solid Phase Extraction (SPE) before being injected into LC-MS/MS. Ice-cold 0.1% formic acid (4 ml) was added in the obtained supernatant to fully protonate the LM species before sample was loaded to the preconditioned SPE cartridges (Strata-X 33 µm polymeric reversed phase, Phenomenex). Once the sample had been totally loaded, cartridges were washed with 0.1% formic acid followed by 15% (v/v) ethanol in water to remove excess salts. Then the LMs from the SPE sorbent bed were eluted by methanol. Solvents were removed using an Eppendorf^®^ 5301 concentrator centrifugal evaporator (Eppendorf). The dried extracts were stored at -80°C immediately for future LC-MS/MS analysis. Prior to analysis the dried extracts were reconstituted in 50 µl methanol, then 10 µl was injected into LC/MS system.

**Sample Profiling:** All components of LC-MS/MS system used are from Shimadzu Scientific Instruments, Inc. The LC system was equipped with four pumps (Pump A/B: LC-30AD, Pump C/D: LC-20ADXR), a SIL-30AC autosampler (AS), and a CTO-30A column oven. The LC separation was conducted on a C8 column (Ultra C8, 150 × 2.1mm, 3 µm, RESTEK). The MS/MS analysis was performed on Shimadzu LCMS-8050 triple quadrupole mass spectrometer. The instrument was operated and optimized under both positive and negative electrospray (+/− ESI) and multiple reaction monitoring modes (MRM). The settings of flow rate and gradient program for the LC system as well as MS/MS conditions are recommended by a software method package for 158 lipid mediators (Shimadzu Scientific Instruments, Inc.,) and further optimized following previously published quantification method [37-39]. The m/z transitions (precursor to product ions) and their tuning voltages were selected based on the best MRM responses from instrumental method optimization software. All analyses and data processing were completed on Shimadzu LabSolutions V5.91 software (Shimadzu Scientific Instruments, Inc., Columbia, MD).
