## Supplemental Figures for "RNA-sequencing reveals a gene expression signature in skeletal muscle of a mouse model of age-associated post-operative functional decline"

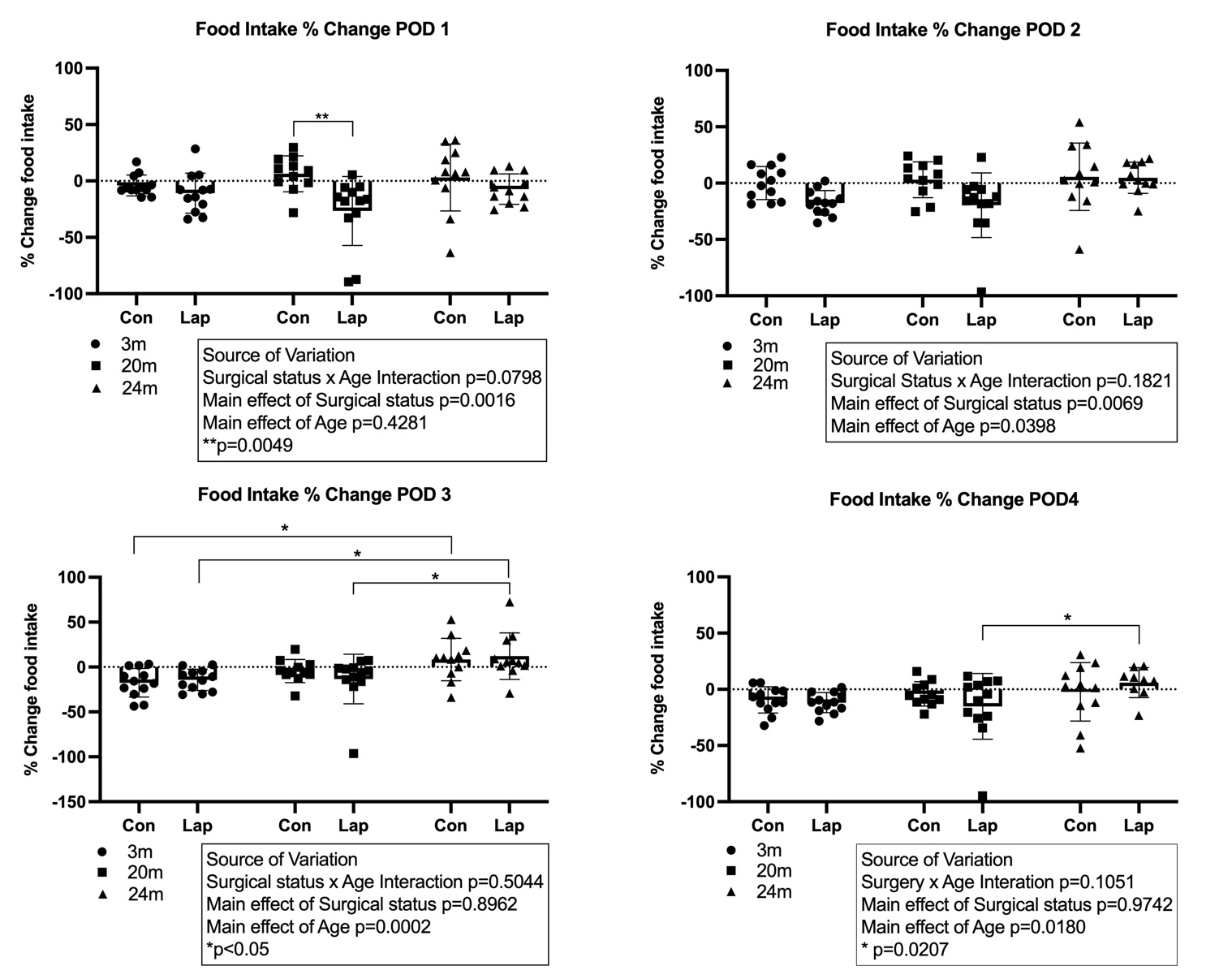

**Supplemental Figure 1: Post-operative changes in food intake.** The percentage change of daily food (rodent chow) consumed compared to the pre-operative food intake was calculated on post-operative days 1-4. Statistical analysis was performed by 2-way ANOVA with Tukey test for multiple comparisons. Error bars denote standard deviation. POD=post-operative day; Con=control; Lap=laparotomy.

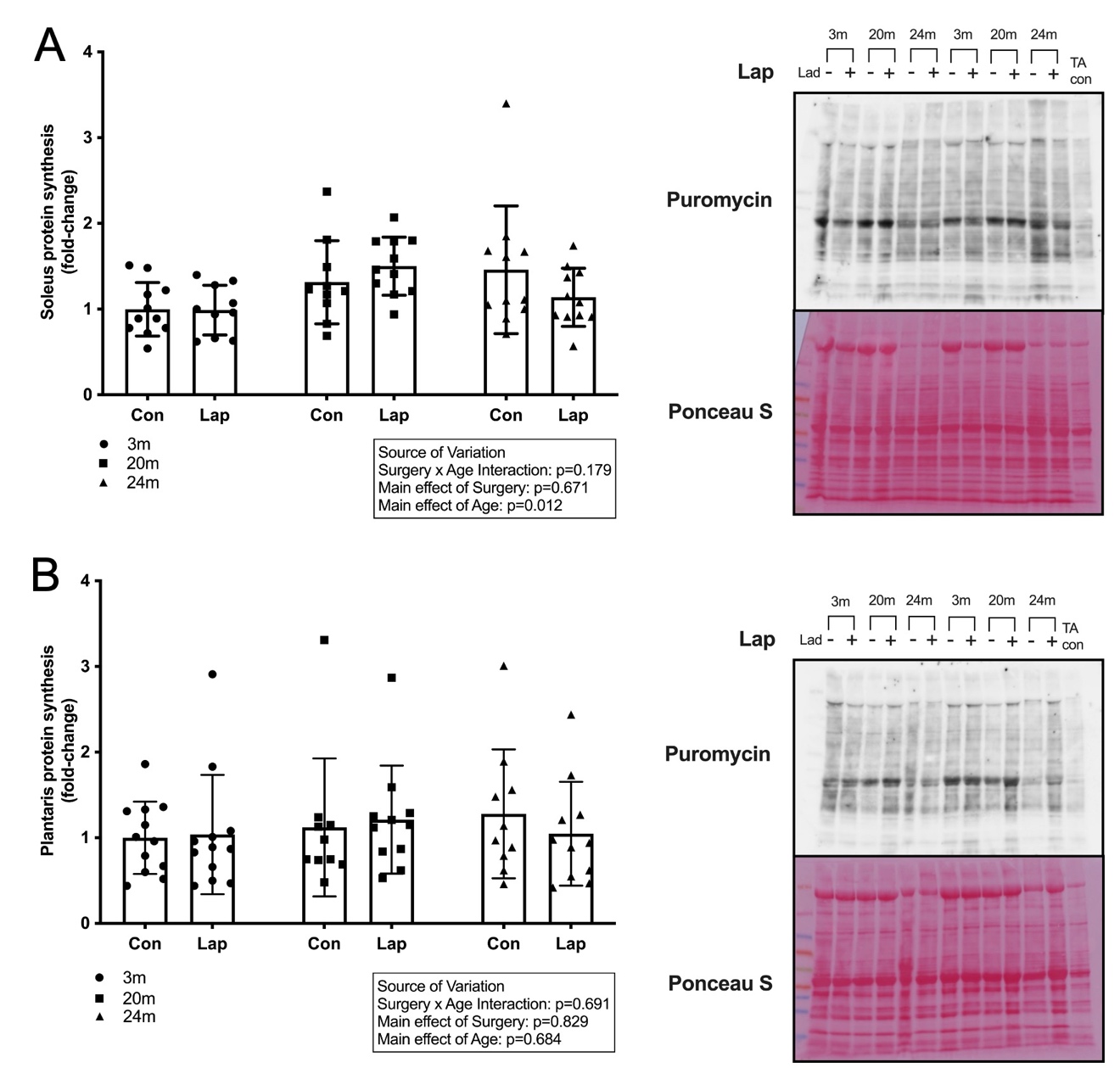

**Supplemental Figure 2: Laparotomy does not affect protein synthesis in soleus or plantaris muscles in C57BL/6N mice.** Quantification of global protein synthesis in soleus (A) and plantaris (B) muscles on post-operative day 5 using surface sensing of translation (SUnSET). N=11-12/group. Statistical analysis was performed by 2-way ANOVA with Tukey test for multiple comparisons. Error bars denote standard deviation. Con=control; Lap=laparotomy.

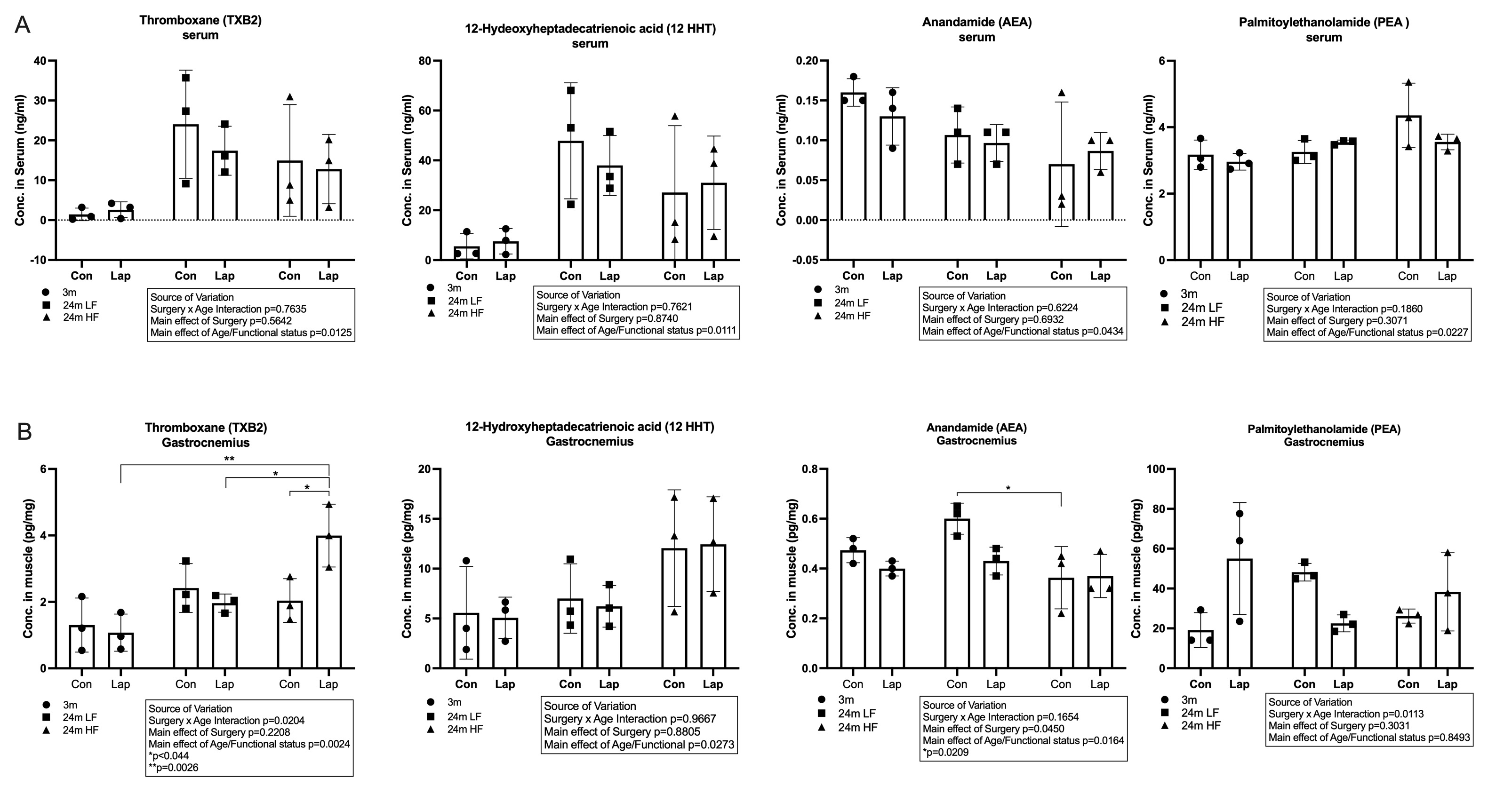

**Supplemental Figure 3: Lipidomics analysis of serum and gastrocnemius muscle showed differences in the main effects of age/functional status in concentration of lipid mediators.** (A) Changes in serum lipid mediator. (B) Changes in gastrocnemius muscle lipid mediators. N=3/group. Statistical analysis performed by 2-way ANOVA with Tukey test for multiple comparisons. Error bars denote the standard deviation. Con=control; Lap=laparotomy; LF=low functioning; HF=high functioning. *Note the y axis scales are not the same due to a large data range.
