## Supplemental Table 1 for "RNA-sequencing reveals a gene expression signature in skeletal muscle of a mouse model of age-associated post-operative functional decline"

| Gene symbol | Gene name | Assay ID |
| --- | --- | --- |
| Ccl2 | chemokine (c-c motif) ligand 2 | Mm00441242_m1 |
| Fox01 | Forkhead box protein 01 | Mm00490672_m1 |
| IL-6 | interleukin 6 | Mm00446190_m1 |
| Mtor | mechanistic target of rapamycin | Mm00444968_m1 |
| Nfkb1 | Nuclear factor of kappa light polypeptide gene enhancer in B cells 1, p105 | Mm00476361_m1 |
| Nfkb2 | Nuclear factor of kappa light polypeptide gene enhancer in B cells 2, p49/p100 | Mm00479807_m1 |
| Ppara | peroxisome proliferator activated receptor alpha | Mm00440939_m1 |
| Pparg | peroxisome proliferator activated receptor gamma | Mm01184322_m1 |
| Ppargc1a | peroxisome proliferator-activated receptor gamma coactivator 1-alpha | Mm01208835_m1 |
| Ppargc1b | peroxisome proliferator-activated receptor gamma coactivator 1-beta | Mm00504720_m1 |
| Rel | reticuloendotheliosis oncogene | Mm01239661_m1 |
| Rela | RELA proto-oncogene, NF-kB subunit | Mm00501346_m1 |
| Relb | avian reticuloendotheliosis viral (v-rel) oncogene related B | Mm00485664_m1 |
| Sln | sarcolipin | Mm00481536_m1 |
| Tuba4a | alpha tubulin | Mm00849767_S1 |
| Ucp-1 | uncoupling protein 1 | Mm01244861_m1 |

**Supplemental Table 1:** Candidate transcriptional regulators in soleus muscles identified via upstream regulator analysis using Ingenuity Pathway Analysis and corresponding TaqMan gene expression assays used for RT-PCR.
